# Faster Cognitive Decline in Dementia due to Alzheimer Disease with Clinically Undiagnosed Lewy Body Disease

**DOI:** 10.1101/510453

**Authors:** TG Beach, M Malek-Ahmadi, E Zamrini, CH Adler, MN Sabbagh, HA Shill, SA Jacobson, CM Belden, RJ Caselli, BK Woodruff, SZ Rapscak, GL Ahern, J Shi, JN Caviness, E Driver-Dunckley, SH Mehta, DR Shprecher, BM Spann, P Tariot, KJ Davis, KE Long, LR Nicholson, A Intorcia, MJ Glass, JE Walker, M Callan, J Curry, B Cutler, J Oliver, R Arce, DG Walker, L-F Lue, GE Serrano, LI Sue, K Chen, EM Reiman

**Affiliations:** Banner Sun Health Research Institute, Sun City, AZ; Banner Alzheimer Institute, Phoenix, AZ; Department of Neurology, Mayo Clinic, Scottsdale, AZ; Cleveland Clinic Lou Ruvo Center for Brain Health, Las Vegas, NV; Barrow Neurological Institute, Phoenix, AZ; Department of Neurology, University of Arizona, Tucson, AZ; Molecular Neuroscience Research Center, Shiga University of Medical Science, Otsu, Japan; Shanghai Green Valley Pharmaceutical; Arizona State University School of Mathematics and Statistics; College of Medicine-Phoenix, University of Arizona; Beijing Normal University

**Keywords:** cognition, autopsy, neuropathology, mental status and dementia tests, diagnosis

## Abstract

Neuropathology has demonstrated a high rate of comorbid pathology in dementia due to Alzheimer’s disease (ADD). The most common major comorbidity is Lewy body disease (LBD), either as dementia with Lewy bodies (AD-DLB) or Alzheimer’s disease with Lewy bodies (AD-LB), the latter representing subjects with ADD and LBD not meeting neuropathological distribution and density thresholds for DLB. Although it has been established that ADD subjects with undifferentiated LBD have a more rapid cognitive decline than those with ADD alone, it is still unknown whether AD-LB subjects, who represent the majority of LBD and approximately one-third of all those with ADD, have a different clinical course. Subjects with dementia included those with “pure” ADD (n = 137), AD-DLB (n = 64) and AD-LB (n = 114), all with two or more complete Mini Mental State Examinations (MMSE) and a full neuropathological examination. Linear mixed models assessing MMSE change showed that the AD-LB group had significantly greater decline compared to the ADD group (β = −0.69, 95% CI: −1.05, −0.33, p<0.001) while the AD-DLB group did not (β = −0.30, 95% CI: −0.73, 0.14, p = 0.18). Of those with AD-DLB and AD-LB, only 66% and 2.1%, respectively, had been diagnosed with LBD at any point during their clinical course. The probable cause of LBD clinical detection failure is the lack of a sufficient set of characteristic core clinical features. Core DLB clinical features were not more common in AD-LB as compared to ADD. Compared with clinically-diagnosed AD-DLB subjects, those that were clinically undetected had significantly lower prevalences of parkinsonism (p = 0.046), visual hallucinations (p = 0.0008) and dream enactment behavior (0.013). Clinical identification of ADD with LBD would allow stratified analyses of ADD clinical trials, potentially improving the probability of trial success.

## Introduction

Dementia due to AD (ADD) is frequently associated with comorbid pathologies that could affect clinical presentation, the rate of clinical progression, and response to investigational or approved treatments. For ADD, there are a host of prevalent co-existing vascular and molecular lesions [1-21].

The presence of comorbid and heterogeneous pathology would be a moot point if it had little or no clinical effect, or if it were only secondarily generated by the primary pathology. But if it resulted in an altered clinical disease progression rate and was not responsive to therapies directed at the “primary” pathology, it could have a significant impact on the response to such therapies. If these were both true, it is apparent that exclusion from ADD clinical trials of subjects harboring such comorbidities would increase the observed effect size and hence clinical trial efficiency.

The most common comorbidity in ADD is Lewy body disease due to aggregation of α-synuclein. Approximately one-half or more of all subjects meeting clinicopathological diagnostic criteria for ADD also have α-synuclein pathology [9;22-24] of the Lewy body type (broadly termed “Lewy body disease”, LBD). Similarly, up to one-half of subjects with dementia and PD (PDD) [25-37] and three-quarters or more of those with DLB, have clinically significant AD pathology [38-41]. In the great majority of subjects with both ADD and LBD, this co-existence is recognized only at autopsy [42-44], currently preventing, except for the minority with clinically-typical DLB, exclusion of LBD subjects from ADD clinical trials. A number of autopsy-validated studies have indicated that cognitive decline is faster in elderly subjects dying with ADD, or any AD neuropathology, who also have undifferentiated LBD [3;45-48] but it is unclear whether or not these results may be driven primarily by DLB subjects with neuropathologically-severe LBD (neocortical stage diffuse Lewy body disease), as disease duration is reportedly shorter in this group [39;49]. The great majority of LBD in ADD subjects does not meet neuropathological diagnostic criteria for DLB, due to insufficient pathology density and brain regional distribution [50-52]. These “AD-LB” cases are most often clinically silent [1] and diagnosed as probable ADD. We have previously reported that AD-LB subjects, as compared to ADD subjects without LBD, depression and Trail-Making Test A scores correlate significantly with LBD pathology but that other neuropsychological variables are not significantly different [53]. This study seeks to determine whether clinical disease progression differs in AD subjects with and without LBD. The prevalences of core DLB clinical features [51;52] were ascertained to help understand the failure to clinically identify many AD-DLB and AD-LB subjects.

## Materials and Methods

### Subject selection

Subjects were selected by database searches of the Banner Sun Health Research Institute Arizona Study of Aging and Neurodegenerative Disorders (AZSAND)/Brain and Body Donation Program (www.brainandbodydonationprogram.org) [54], a subset of whom were also enrolled in the National Institute on Aging Arizona Alzheimer’s Disease Core Center. Search criteria specified that subjects died with dementia, two or more complete Mini Mental State Examinations (MMSE) and a full neuropathological examination after death. Of these, selected subjects met “intermediate” or “high” National Institute on Aging-Reagan Institute (NIA-RI) clinicopathological criteria [55] for ADD, with or without also meeting “intermediate” or “high” clinicopathological criteria for DLB [51;52], or alternatively, for a group termed as AD-LB [50], also having pathologically-confirmed CNS LBD but not meeting DLB pathology distribution and density thresholds. For all subjects, other major neuropathological disorders were excluded, including cases clinicopathologically defined as vascular dementia, hippocampal sclerosis, frontotemporal lobar degeneration with TDP-43 proteinopathy, Pick’s disease, progressive supranuclear palsy, multiple system atrophy, corticobasal degeneration, Huntington’s disease, large acute cerebral infarcts, motor neuron disease and primary or metastatic malignant brain tumors. Selected subjects (Table 1) were divided into three groups: ADD (n = 137), AD-DLB (n = 64) and AD-LB (n =114).

**Table 1.**
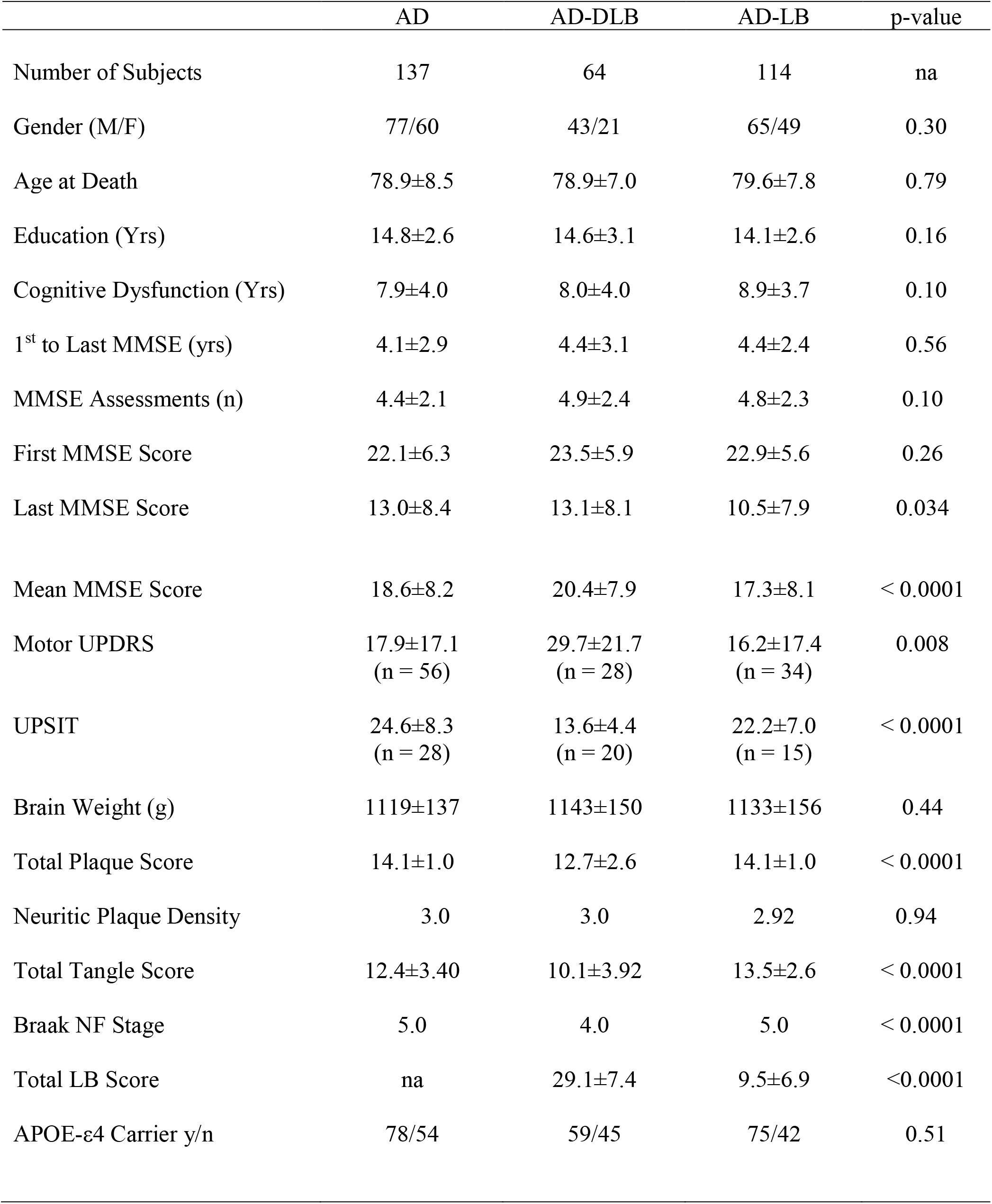
Demographic and post-mortem characteristics of study subjects. Means and standard deviations are shown; medians are shown for neuritic plaque density and Braak stage. Cognitive dysfunction duration is the number of years elapsed between first clinical appearance and death. Motor UPDRS scores for all groups were done off medications; the score shown is the final score prior to death. Group comparisons are by one-way analysis of variance or Kruskall-Wallis analysis of variance, as appropriate. Other comparisons are with chi-square tests or unpaired, 2-tailed Student t-tests. MMSE = Mini Mental State Examination; UPDRS = Unified Parkinson’s Disease Rating Scale; UPSIT = University of Pennsylvania Smell Identification Test; NF = neurofibrillary; LB = Lewy body; ApoE-E4 = apolipoprotein E, ε4 allele

### Subject characterization

Most subjects had serial standardized research cognitive evaluations, done by teams of nurses, medical assistants, behavioral neurologists, neuropsychologists and psychometrists using standardized research-quality assessment batteries [54], including the National Alzheimer’s Coordinating Center (NACC) Uniform Data Set (UDS). To estimate group prevalences of DLB core clinical features [51;52], the presence or absence of parkinsonism, visual hallucinations, Mayo Sleep Questionnaire findings [56-59] and/or clinical history consistent with REM sleep behavior disorder (RBD), as well as fluctuations in attention or cognition were recorded for each subject. A subset (n = 63) received, every third year, a University of Pennsylvania Smell Identification Test (UPSIT) [60-62]. Another subset of subjects (58 AD, 31 AD-DLB and 36 AD-LB) additionally received serial assessments, including the Unified Parkinson’s Disease Rating Scale (UPRDS), administered by subspecialist movement disorders neurologists. The presence of PD, parkinsonism and hallucinations was additionally noted by standardized medical history questionnaires, and by review of private medical records.

All subjects received identical neuropathological examinations, including summary brain density measures for total amyloid plaques, neurofibrillary tangles, Lewy body pathology regional and summary density scoring, and staging using the Unified system [50], as well as assignment of CERAD neuritic plaque density and Braak neurofibrillary stage, as described previously [54].

### Statistical analysis

Demographic and post-mortem characteristics were analyzed using one-way analysis of variance (ANOVA), Kruskall-Wallis analysis of variance, Chi-square tests and t-tests as appropriate. Linear mixed-effects models were used to assess MMSE slope differences among the three groups. The first model included fixed effects for age (mean-centered), baseline MMSE score, diagnostic group, a visit age by group interaction term as covariates, and the random slopes for each subject. The second model added education, sex, and AD neuropathological severity (intermediate or high) according to the NIA-RI classification [55]. Fitted MMSE values from the second model were used to calculate annualized change for each of the groups.

## Results

Clinical, demographic and post-mortem characteristics of the compared groups are shown in Table 1. No significant differences were noted for gender, age at death, proportion of subjects carrying the apolipoprotein E ε4 allele, education, duration of cognitive dysfunction until death, elapsed time between first and last MMSE, mean number of MMSE tests, duration between first assessment and death, or post-mortem interval. The groups were not significantly different on their first MMSE scores (p = 0.26) but differed on their mean (p < 0.0001) and final (p = 0.034) scores, with the AD-LB group having the lowest scores. Motor scores on the UPDRS and scores on the UPSIT differed significantly across groups (for both, p < 0.0001), driven by greater impairment in the AD-DLB group. Most subjects had a dementia diagnosis at study entry but approximately 20-35% were originally classified as cognitively normal or having mild cognitive impairment (MCI); the proportions were not significantly different between groups. Neuropathologically, subjects did not differ in their brain weights or neuritic plaque densities but were significantly different in total plaque scores, total tangle scores and Braak stage, with the AD-DLB group consistently having the lowest AD pathology scores. As expected by group definition, Lewy body pathology brain load was much higher for the AD-DLB group as compared to the AD-LB group. The lower AD-related pathology scores for the AD-DLB group were also expected due to classification rules for DLB that are more permissive when lower AD pathology densities are present [51].

The results from the first linear mixed-effects model showed that the AD-LB group had significantly greater decline compared to the ADD group (β = −0.69, 95% CI: −1.05, −0.33, p<0.001) while the AD-DLB group did not (β = −0.30, 95% CI: −0.73, 0.14, p = 0.18). Results from the second model with the additional covariates yielded nearly identical results: AD-LB (β = −0.68, 95% CI: −1.04, −0.32, p<0.001); AD-DLB (β = −0.29, 95% CI: −0.72, 0.14, p = 0.18). The MMSE slopes for the AD-DLB and AD-LB groups were not significantly different in either model (for both, p = 0.17). Linear mixed model-derived annualized MMSE changes for each group are shown in Figure 1. The AD-LB group showed an annualized MMSE change of −2.16 ± 0.76 points while for the AD-DLB and ADD groups, these were −1.77 ± 0.86 and −1.48 ± 0.70, respectively (Table 2).

**Figure 1.**
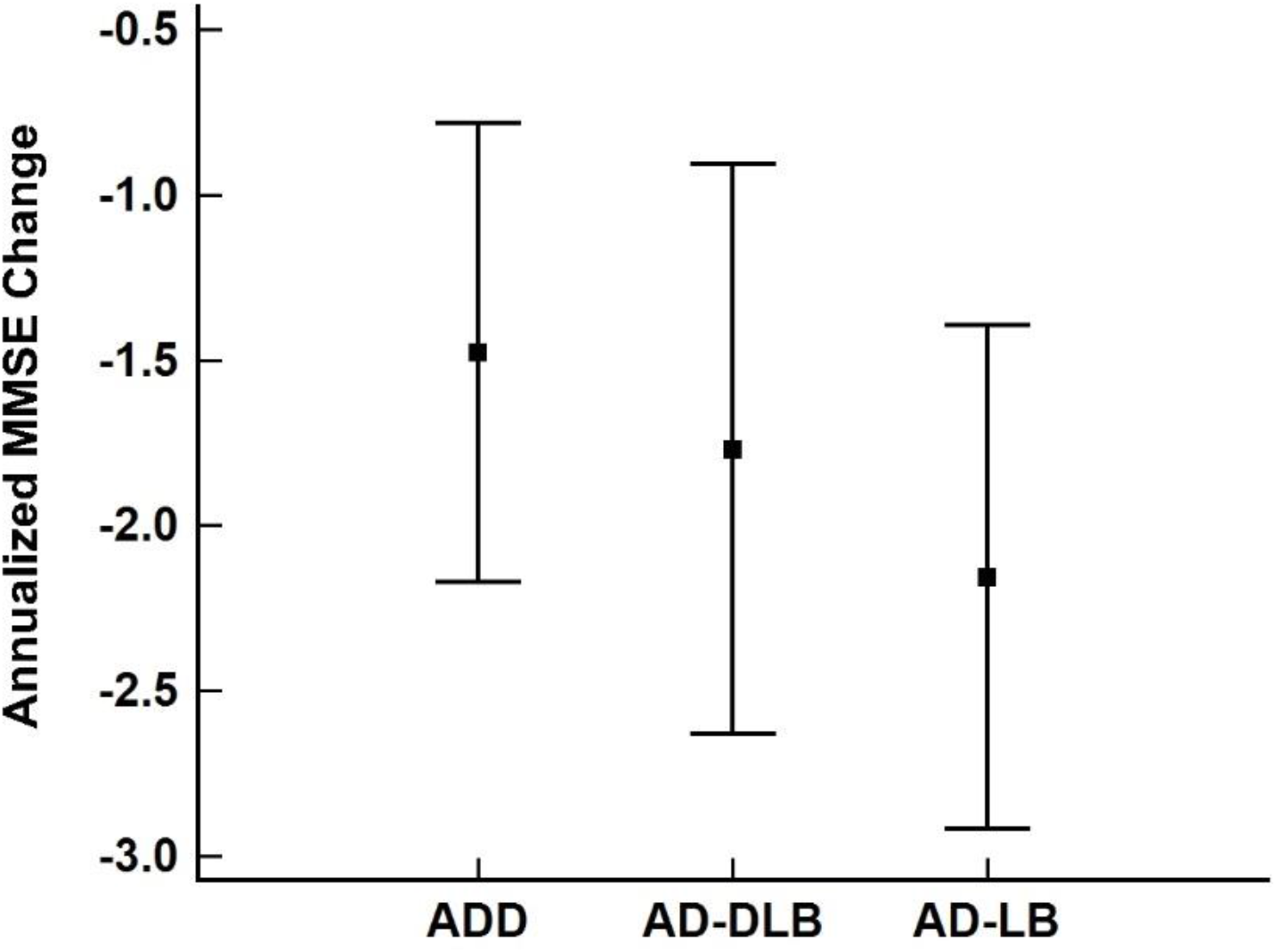
Annualized MMSE change group differences. Means and standard deviations are shown.

**Table 2.**
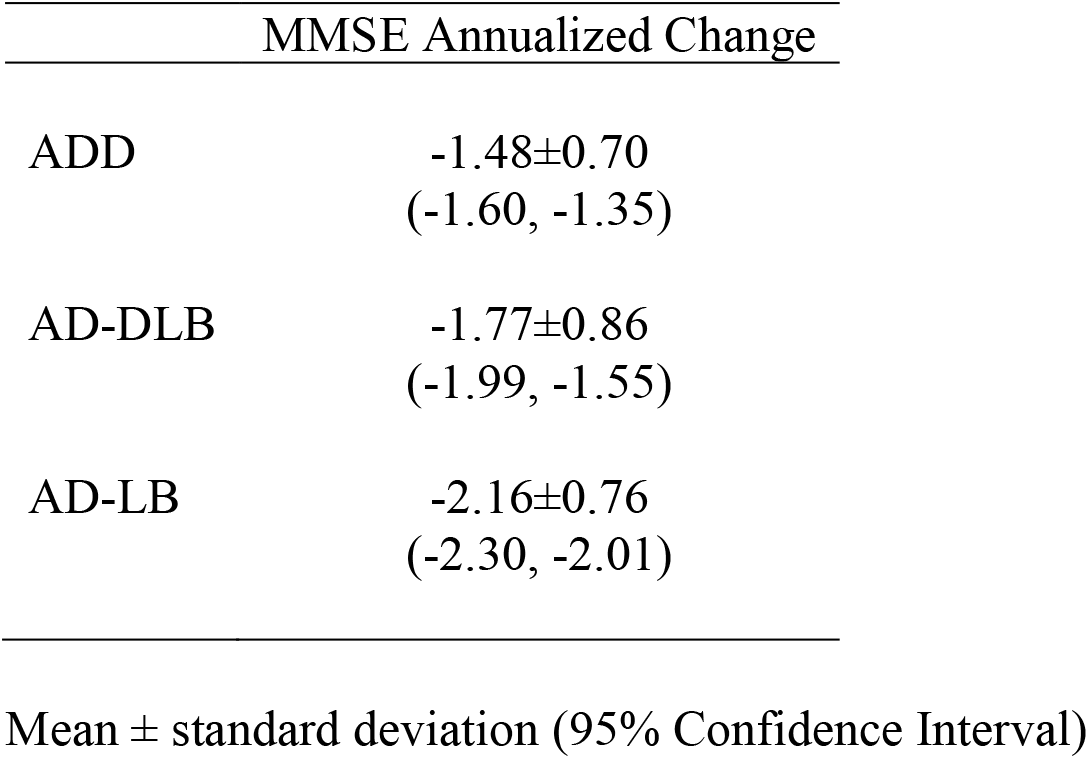
Annualized MMSE change for ADD, AD-DLB, and AD-LB groups.

Lewy body disease diagnosed at autopsy was often unrecognized during life, even when examiners were specialized behavioral neurologists, neuropsychologists and psychometrists using standardized research-quality assessments, including the NACC UDS, UPDRS and Mayo Sleep Questionnaire. Of those meeting both NIA-RI intermediate or high neuropathological criteria for AD as well as DLB III [51] neuropathological criteria for intermediate or high likelihood for DLB, and that had a final postmortem (blinded to neuropathology) clinical diagnostic conference from the specialist research team, 21/32 (66%) had been clinically diagnosed at least once (any Lewy body related clinical diagnosis at any time during their clinical course was accepted) with DLB, “Lewy body dementia” or “Lewy body variant of Alzheimer’s disease” by either the research teams of specialized neurologists and neuropsychologists or by private medical practitioners. However, only 15/32 (46.9%) of these were diagnosed as DLB at the specialist research team’s final postmortem (blinded to neuropathology results) clinical diagnostic conference. For those with AD-LB, the percentage diagnosed by the research teams at any time with a Lewy body related diagnosis was 2/93 (2.1%). In comparison, for those with AD and no postmortem evidence of Lewy body disease, 4/85 (1.6%) had been diagnosed at least once as having a Lewy body related diagnosis.

To determine why the diagnosis of LBD was being missed relatively frequently by the clinical research teams, we examined the group prevalences of core DLB clinical features, including parkinsonism, visual hallucinations, cognitive fluctuations and history consistent with RBD and/or dream enactment behavior (Tables 3 and 4). Of the neuropathologically-defined AD-DLB cases evaluated at least once by the specialized clinical research teams, 58% had parkinsonism, 47% had hallucinations, 33% had cognitive fluctuations and 18% had a history of RBD and/or dream enactment behavior. Of the AD-LB subjects that had at least one evaluation by the research teams, the core features were much less common: 30% had parkinsonism, 11% had hallucinations, 13% had fluctuations and 5% had RBD and/or dream enactment behavior. The difference in prevalences were all significantly greater for AD-DLB versus the AD-LB and ADD groups (p < 0.05) (Table 3). Emphasizing the relative infrequency of core features in the AD-LB group, their prevalences did not significantly differ from the ADD group. The presence of all core features, except cognitive fluctuation, were significantly more common in the clinically-recognized AD-DLB subjects as compared to those meeting DLB neuropathological criteria (DLB Consortium intermediate or high) but who were unsuspected as such during life (Table 4).

**Table 3.** Prevalence of DLB core clinical features in the three groups. Only data from subjects seen at least once by clinical research teams is considered. RBD/DEB = clinical diagnosis of REM sleep behavior disorder or presence of dream enactment behavior.

|  | parkinsonism* | hallucinations* | fluctuations* | RBD/DEB* |
| --- | --- | --- | --- | --- |
| AD-DLB | 32/55 (58%) | 26/55 (47%) | 18/55 (33%) | 10/55 (18%) |
| AD-LB | 26/93 (30%) | 10/93 (11%) | 13/93 (14%) | 5/93 (5%) |
| ADD | 28/109 (26%) | 17/109 (16%) | 22/109 (20%) | 5/109 (5%) |

\* = chi-square $p < 0.05$

**Table 4.** Prevalence of DLB core features in AD-DLB subjects with and without a clinical DLB diagnosis. Only data from subjects seen at least once by clinical research teams is considered. Clin/path = subjects diagnosed as DLB both clinically and neuropathologically. Path = subjects without a clinical diagnosis of DLB but neuropathologically meeting DLB III intermediate or high criteria. RBD/DEB = clinical diagnosis of REM sleep behavior disorder or presence of dream enactment behavior.

|  | parkinsonism* | hallucinations* | fluctuations | RBD/DEB* |
| --- | --- | --- | --- | --- |
| DLB<br>(clin/path) | 18/24 (75%) | 16/24 (67%) | 9/24 (38%) | 8/24 (33%) |
| DLB<br>(path) | 14/31 (45%) | 10/31 (32%) | 9/31 (29%) | 2/31 (6%) |

\* = $p < 0.05$

## Discussion

The results confirm prior findings by others that cognitive decline is faster in subjects with autopsy-confirmed ADD if they also have co-existing LBD neuropathology [3;39;45-49;63]. A new finding is that this increased rate of decline is restricted to AD-LB subjects who do not meet neuropathological density and distribution criteria for DLB. A large fraction of AD-DLB cases, and virtually all the AD-LB cases, were unrecognized as having LBD during life, even though they had at least one and in many cases annual evaluations by specialized research teams including both behavioral and movement disorders subspecialist neurologists. As AD-LB is much more common than AD-DLB (33% and 16% of all ADD cases, respectively, our unpublished data), this means that approximately 40% or more of ADD subjects have co-existing, clinically-silent LBD that accelerates their disease progression. Aside from complicating clinical trial analyses using rate of cognitive decline as a marker of therapeutic response, it is still unknown whether or not therapy directed at AD molecular targets will have any effect on LBD. It is possible that an agent might slow or halt the growth of AD pathology but be clinically ineffective due to the unobstructed advance of LBD.

It is surprising that LBD conferred a faster rate of cognitive deterioration regardless of the severity and extent of LBD pathology. The difference from ADD was in fact restricted to the AD-LB group, despite the greater LBD pathology densities and regional brain distribution in AD-DLB. In AD-DLB subjects, LBD pathology has most often spread to the neocortical stage, while in AD-LB it is most often restricted to a limbic-predominant stage [50], and the density of LBD pathology is almost always greater in AD-DLB, in every involved brain region. As the difference in rate of decline between AD-LB and ADD is maintained even after adjustment for level of AD neuropathology, it seems likely that AD-LB is an inherently more aggressive disease than ADD, while AD-DLB is similar to ADD.

It has been reported that AD-DLB may be a more malignant condition than ADD, based on a reportedly more rapid disease course from first presentation to nursing home placement or death [64;65]; conversely, the rate of cognitive decline has most often been found to be similar in the two conditions [64-66]. Others have shown that survival time, in those with clinically probable, neuropathologically-confirmed DLB, is significantly shorter with diffuse neocortical LBD as compared with LBD restricted to the limbic regions [49] and that this is at least partially related to greater AD pathology, the presence of lacunar infarcts and parkinsonism. This cited study did not compare survival between ADD and AD-DLB. In the current study, age at death and survival did not differ between ADD and AD-DLB, even when AD-DLB cases were restricted to those with neocortical stage (results for survival and for the separated neocortical group not shown). Differences between our study and that cited above [49], aside from lack of an ADD comparison group in the latter, include our case selection, based on neuropathologically-defined DLB, rather than a combined clinical and neuropathological definition, as well as our exclusion of cases meeting vascular dementia neuropathological criteria, resulting in an almost complete lack of lacunar infarcts in our AD-DLB group (only 3 cases had any lacunar infarcts). One other study [39], based on neuropathologically-defined DLB, confirmed a shorter disease duration to death in neocortical stage LBD as compared to limbic stage, but did not include an ADD comparison group.

A limitation to these conclusions is that the MMSE is a single, relatively simple test and comprehensive neuropsychological test comparisons would be more informative. Unfortunately, more sophisticated neuropsychological tests often cannot be performed in subjects with advanced dementia and so the sample size is much reduced. In a reduced subject set with full neuropsychological test battery results, we previously reported that AD-LB and ADD subjects differed only on depression and Trail-Making A scores [53].

Whether or not LBD initiation or spread is independent, in ADD subjects, of the AD neuropathology, is not yet established. While experimental models have shown potential synergistic or overlapping molecular mechanisms generating AD and LBD pathology [67-70], it is also apparent that LBD is commonly present in subjects where AD pathology is clearly primary, such as in autosomal dominant mutations of PS1 and APP [70-75], or even in other cerebral amyloidoses [76]. It is possible, then, that much of the LBD in AD subjects is produced as a secondary event and thus effective anti-amyloid or anti-tau agents might also be effective against LBD. To investigate the possibility of differential effects of therapeutic agents on AD and LBD pathology, it will be necessary to first acquire much more sensitive and specific LBD clinical diagnostic methods.

Clinically-typical DLB is readily and accurately identified when the core clinical features are present but in this set of subjects these were only 31% of all those with ADD and LBD. The cognitive presentation may be helpful, as initial impairment in attention, executive function and/or visuospatial function may be more common than in AD without LBD [77]. The presence of REM sleep behavior disorder (RBD) is a strong predictor of underlying LBD [78] but this relationship is weaker for the relatively low LBD pathology loads typical of AD-LB [56;79]. In this study, as in others [80;81] decreased olfaction is strongly associated with LBD. The association was restricted to the AD-DLB group however, as UPSIT scores for the AD-LB and pure AD groups were not significantly different. Although the sample size for all groups is relatively small, these results suggest that hyposmia or anosmia may be a key diagnostic feature of DLB.

Useful laboratory-based biomarkers for LBD are not yet available. Functional dopamine imaging (e.g. DaTscan) and myocardial scintigraphy with [123I]meta-iodobenzylguanidine (MIBG) have both been used as diagnostic adjuncts for DLB [82;83] with promising but not yet definitive results from small autopsy-confirmed studies [84]. As both nigrostriatal degeneration and peripheral α-synuclein pathology are largely absent in AD-LB [50;85;86], these approaches may prove to be of use as biomarkers for AD-DLB but would likely not be of value for distinguishing AD-LB from ADD. Biofluids and PET imaging approaches have so far been unsuccessful in providing the required accuracy for identifying LBD [87-89]. Simulation studies have suggested that cortical biopsy [90-93] would have high sensitivity and specificity for DLB, and usage of needle cores rather than open biopsy may reduce morbidity to acceptable levels [90] but would not detect AD-LB subjects as they mostly lack cortical LB pathology. Biopsy of the peripheral nervous system [94], particularly the submandibular gland [86;95-97], shows promise for diagnosing DLB but peripheral LBD pathology is relatively rare in AD-LB. Autopsy studies have suggested that biopsy of the olfactory bulb would identify more than 90% of all subjects with LBD [98]. Better clinical diagnostic methods for LBD are critically needed, as the identification of AD-LB subjects entered into ADD clinical trials would enable their exclusion or a stratified analysis that would potentially increase the probability of trial success.

## Acknowledgements

The Brain and Body Donation Program has been supported by the National Institute of Neurological Disorders and Stroke (U24 NS072026 National Brain and Tissue Resource for Parkinson’s Disease and Related Disorders), the National Institute on Aging (P30 AG19610 Arizona Alzheimer’s Disease Core Center), the Arizona Department of Health Services (contract 211002, Arizona Alzheimer’s Research Center), the Arizona Biomedical Research Commission (contracts 4001, 0011, 05-901 and 1001 to the Arizona Parkinson’s Disease Consortium) and the Michael J. Fox Foundation for Parkinson’s Research.

